# Ex-STORM: Expansion Single Molecule Super-resolution Microscopy

**DOI:** 10.1101/049403

**Authors:** Zhisong Tong, Paolo Beuzer, Qing Ye, Josh Axelrod, Zhenmin Hong, Hu Cang

**Author notes:** These authors contribute equally to this manuscript (Zhisong Tong, Paolo Beuzer).

## Abstract

We present critical improvements to recently invented Expansion Microscopy (ExM), which resolve the incompatibility between ExM and single molecule super-resolution microscopy STORM. Specifically, the improved ExM circumvents the massive, 50-100%, bleaching of fluorophores during sample preparations, and preserve the efficiency of enzymatic oxygen scavenging systems in expanded samples. These improvements open up new avenues for Ex-STORM - expanding a sample mechanically, and then visualizing the sample with STORM.

## INTRODUCTION

Recently, Chen^1^ *et al.* demonstrated a novel approach to overcome the long-standing challenge of diffraction limit in light microscopy. Rather than improving the optics of a microscope^2, 3^ or using photo-controllable fluorescent molecules^4, 5^, they proposed to expand specimens mechanically to magnify sub-diffraction-limit features. To ensure that the expansion would be isotropic, they developed a method to imprint a specimen’s features of interest onto a polyacrylamide hydrogel, and then expand the polyacrylamide hydrogel homogeneously by 3 to 4 folds (Fig. 1a). This method, termed Expansion Microscopy, or ExM, could enhance the lateral resolution of a confocal microscope to ~60 nm^1^.

**Fig. 1.**
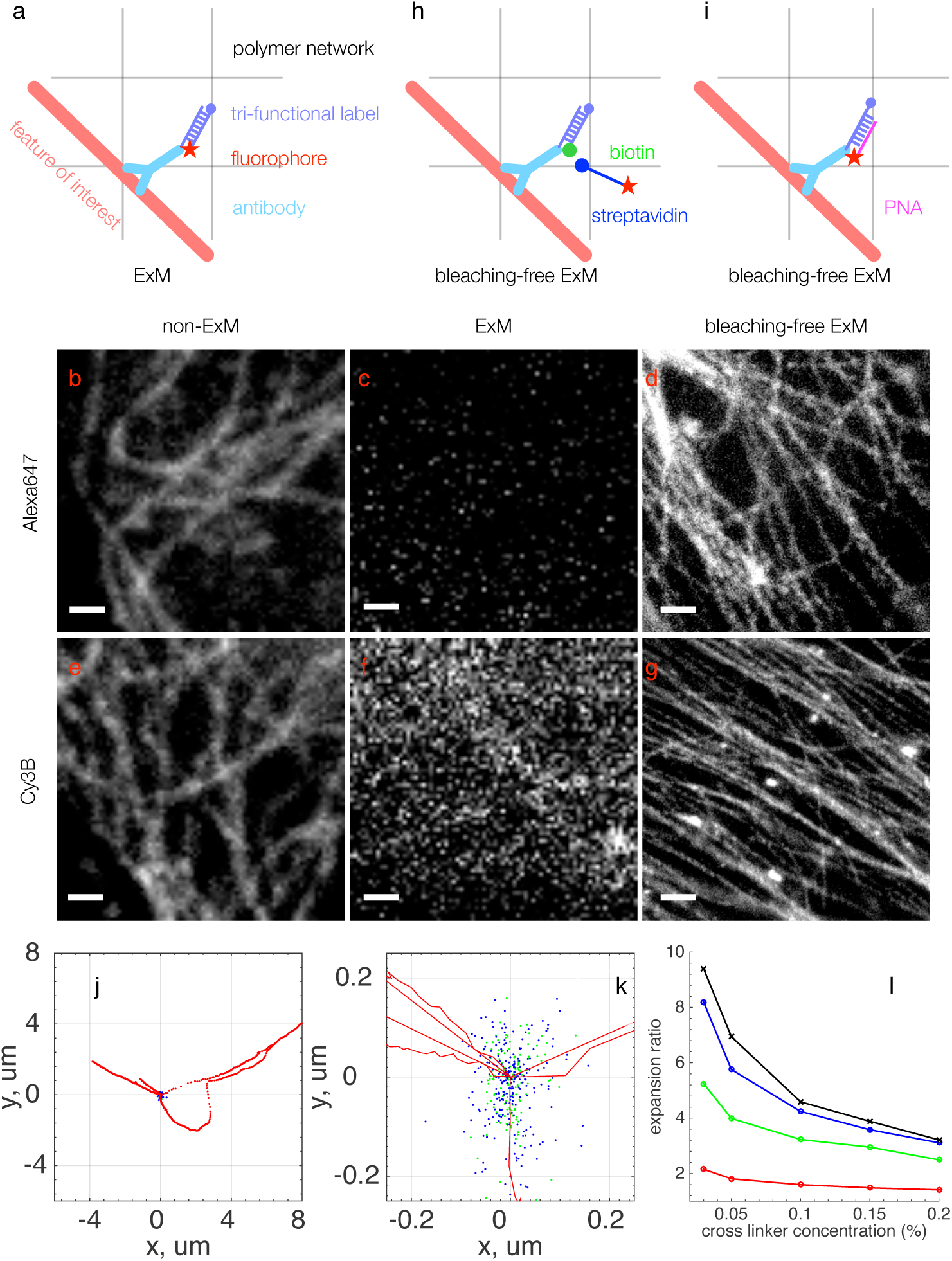
Bleaching Free ExM. (a). ExM imprints features of interest to an expandable polymer hydrogel network by using oligonucleotide conjugated antibodies and complementary tri-functional labeled oligonucleotides. The polymerization of the polymer network bleaches fluorophores massively. Epi-fluorescence images of microtubules stained with Alexa647 (b-d) and Cy3B (e-g). Microtubules are completely disappeared in ExM samples with Alexa647 dyes (c), and are barely visible with Cy3B dyes (f). Our bleaching-free ExM lose no signals at all for both dyes (d and g). The scale bars in the images are 1μm. The magnification in (d) and (g) has been normalized to account for the expansion of the gel. The high background in (f) is due to the adjustment of image contrast in order to visualize the faint microtubule fibers. (h). A bleaching-free ExM using a biotin-conjugated tri-functional label. (i). A bleaching-free ExM that allows multi-color labeling by crosslinking the oligonucleotides tagged on the antibodies directly to the polymer network; complementary PNA probes stain the oligonucleotides after the polymerization. (j). The trajectory of 40 nm red fluorescence beads in an expanded hydrogel on a normal cover glass (red) shows significant drifting of the hydrogel in a period of 8 minute, while both the trajectories of beads in similar gels on poly-lysine coated cover glass (green) or embedded in agarose gel (blue) exhibit less than 300 nm drifting in 15 minutes. (k). A zoomed in view of (j). (l) The expansion ratio *vs* the cross linker concentration of hydrogels are characterized in 4 buffers. The 4 curves correspond to ddH2O (black), diluted STORM imaging buffer (1 mM tris pH8.0 and 10 mM glucose, blue), normal STORM imaging buffer (10 mM tris pH8.0, 10% glucose, and 1% β-mercaptoethanol, green), and proteinase K digestion buffer (50 mM Tris pH8, 1 mM EDTA, 0.5% Triton X-100, 0.8 M guanidine HCl, and 2.9 M NaCl, red).

Higher spatial resolution could be achieved, if one can integrate ExM with Stochastic Optical Reconstruction Microscopy (STORM/*direct*STORM) - expanding a sample mechanically to magnify the features of interest first, and then visualizing the sample with STORM.

However current ExM procedure is not compatible with STORM^1^. Firstly, fluorescence dyes are massively bleached during sample preparation^1^. The bleaching is particularly severe for some of the most used fluorescence dyes. For examples, Alexa647/Cy5 and Cy3B are two of best photo-switching dyes screened out by Zhuang’s group from 26 common fluorescence dyes for STORM^6^; Fig. 1b to g show that, using microtubules in U2OS cells as a demonstration, nearly 100% of Alexa647 and Cy3B molecules are bleached in the ExM samples (Fig. 1c and f). Since insufficient fluorescence dyes can induce serious artifacts to STORM images^7^, performing STORM on expanded samples is difficult.

Secondly, the sample immobilization scheme of ExM significantly reduces the efficiency of enzymatic oxygen scavenging systems that are critical to maintain reliable photo switching for STORM. Currently, ExM immobilizes the expanded polyacrylamide hydrogel on a cover glass for imaging by embedding the expanded polyacrylamide hydrogel in another polyacrylamide or agarose gel^1^; enzymatic oxygen scavenging systems perform poorly in this embedding hydrogel. As a result, significantly fewer blinking events are observed, reducing the resolution of reconstructed STORM images.

We report here two novel expansion procedures that circumvent the bleaching of fluorophore by inducing the fluorphore after the polymerization step that causes the massive bleaching. While 50-100% of fluorophores are bleached in the original ExM^1^, bleaching is completely prevented with our methods, which allows imaging of single molecules in expanded samples. In addition, we develop a new method to immobilize expanded polyacrylamide hydrogel on cover glass for STORM without the embedding hydrogels; this immobilization method is fully compatible with the oxygen scavenging systems. Taken together, our improvements enable synergetic integration of ExM with STORM. 2.5 folds increase of the localization precision from conventional STORM is achieved, demonstrating the potentials of Ex-STORM.

## RESULTS AND DISCUSSION

### Bleaching Free ExM

Fig. 1a illustrates the procedure of original ExM. Fixed and permealized cells are stained with antibodies conjugated with a short, single strand DNA oligonucleotide. The DNA oligonucleotide is then hybridized to a tri-functional label, comprising a methacryloyl group at the 5’ end, capable of participating in polymer network polymerization, a fluorophore at 3’ end, and a single strand oligonucleotide complementary to the sequence conjugated on the antibodies. The cells are then immersed into an acrylamide monomer solution of high ionic strength. Polymerization is then initiated by ammonium persulfate (APS) with tetra-methyl-ethyl-enediamine (TEMED). The tri-functional labels are covalently linked to the polymer network during the polymerization. After the antibodies are digested by protease K, dissociating the tri-functional labels from the specimens, the polyacrylamide hydrogel is expanded by dialysis in a low ionic strength buffer. The radicals generated during the polymerization step are responsible for the massive bleaching^1^.

The first strategy to overcome the bleaching exploits biotin/streptavidin binding (Fig. 1h). We replace the 3’ fluorophore modification in the original ExM tri-functional label with a biotin. After the polymerization, when the free radicals are consumed, the polymer gel is then stained with Alexa647 or Cy3B conjugated streptavidin. In this way, fluorescent dyes avoid exposure to the radicals that cause the massive bleaching.

We develop a second strategy that could be scaled for multi-color imaging by using peptide nucleic acid (PNA), an artificially synthesized polymer similar to DNA. Fig. 1i illustrates this strategy. We conjugate a, 12 bp long, single strand DNA oligonucleotide to secondary antibodies. The 3’ end of the oligonucleotide is conjugated to the antibody, and 5’ end comprise a methacryloyl group. During the polymerization step, the oligonucleotides were covalently cross-linked to the polymer hydrogel network through the methacryloly group. After the steps of proteinase K digestion and expansion of the polymer hydrogel, the single strand DNA oligonucleotides anchored on the polyacrylamide hydrogel could be post-stained by complementary PNA probes tagged with Cy3B or Alexa647 fluorescence dyes. This approach is similar to Fluorescence in situ hybridization (FISH). One can achieve multi-color labelling^1^, by choosing different combinations of oligonucleotide and PNA sequences. Three combinations are provided in the Materials and Methods sections.

Because the polyacrylamide hydrogel traps unconjugated fluorophores, to wash them out would take a few days; we developed an efficient washing procedure based on electrophoresis to wash away the unconjugated fluorophores for single-molecule imaging. The polyacrylamide hydrogel is embedded in a 0.8% agarose gel placed in a horizontal gel box filled with 1X Tris/Borate/EDTA buffer (TBE). The electrophoresis is run at 100V. We use a fluorescence gel imaging system (Syngene Pxi6) to monitor the level of residual unconjugated fluorophore in the sample. The electrophoresis is continued until the unbound dye is not detectable, which typically takes about 45-60 minutes. After retrieving from the agarose gel, the sample is expanded following the standard ExM procedure^1^.

Fig. 1d and g show that the post staining completely prevents bleaching; the fluorescence intensity of the expanded samples post stained with Alexa647 (Fig. 1d) and Cy3B (Fig. 1g) reaches a similar level to the intensity of non-expanded samples (Fig. 1b and e).

### Immobilizing Expanded Hydrogel For STORM Without Compromising The Efficiency Of Oxygen Scavenging System

Another challenge to achieve Ex-STORM is to maintain the efficiency of oxygen scavenging system^6^. The oxygen scavenging system, (40mM D-glucose, 0.5 mg/ml glucose oxidase (Sigma G6125), 40 μg/ml catalase (Sigma C1345), and 143 mM β-mercaptoethanol (Sigma M6250)), critically determine the quality of reconstructed STORM images. In ExM, hydrogel samples are embedded in agarose gel for stabilization during imaging^1^, however agarose gel reduces the efficiency of the oxygen scavenging systems, and causes a reduction in blinking events and poor image quality. We found that the expanded polyacrylamide hydrogel bound strongly to poly-lysine treated cover glass, (cover glasses incubated in 0.1% poly-lysine solutions for 10 minutes), possibly through strong Coulomb interactions between positive charged glass surface and negative charged poly-acrylate. Fig. 1j-k shows that while the expanded polyacrylamide hydrogel drift about 8 μm in 8 minutes on non-coated glass, out of the camera’s field of view, both the hydrogel on a poly-lysine coated glass and embedded in agarose gel drift less than 300 nm in 15 minutes. The 300-nm level drifting, which also includes thermal drift from microscope stages and sample holder, can be corrected by cross correlations images analysis and fiduciary markers. Therefore the expanded polyacrylamide hydrogel can be immobilized in STORM imaging buffer (10 mM tris pH8.0, 10% glucose, and 1% β-mercaptoethanol) for long-term image acquisitions, without the need of an embedding hydrogel that reduces the efficiency of oxygen scavenging system.

### Ex-STORM Increases The Localization Precision By 2.5 Folds From Conventional STORM

The expansion ratio of hydrogel depends on the ionic strength of the buffered solutions (Fig. 1l). The maximum expansion is achieved in pure water. For STORM experiments, hydrogel samples are expanded directly in STORM imaging buffer. Lower cross linker (N,N’-Methylenebis(acrylamide)) concentrations leads to higher expansion factors, however gels become fragile and difficult to manipulate. Dilution of the STORM imaging buffer by 10 folds can achieve higher, 4 fold, expansion ratio; however, the activity of the oxygen systems drops quickly in low capacity buffer^8^. Therefore hydrogel of 0.15% cross linker concentration, which expands ~3.1 folds in STORM imaging buffer, is used for most experiments.

Expanding a sample by 3 folds increases the localization precision of Ex-STORM by 2.5 folds from conventional STORM. Following ref 9, we characterize the localization precision of Ex-STORM with individual AlexaFluor-647 molecules scattered in expanded hydrogels (Fig. 2). The gel expands 3.1 times. The standard deviation of multiple localizations of the same molecule, *σ*, is used as a metric to determine the localization precision; they are 3.4, 3.7, and 9.6 nm in *x, y*, and *z* direction, respectively, representing 2.5 folds enhancement, on average, from conventional STORM, (8.0, 10.1 and 29.6 nm, respectively) (Fig. 2a-d). The resolutions of the corresponding Ex-STORM images in *x, y* and *z* directions, measured in full-width-half-maxima (FWHM), are 8.0, 8.7, and 22.6 nm respectively; 2.5 folds higher than conventional STORM (18.8, 23.8, and 69.7 nm respectively, which are similar to or better than values reported^9^). The sub 10-nm lateral and 20-nm axial resolution has only been previously achieved by using dual-objective STORM^9^, which requires complex 4-pi configuration and is not available to most laboratories. In addition, the improved, bleaching free ExM is fully compatible with the dual-objective STORM.

**Fig. 2.**
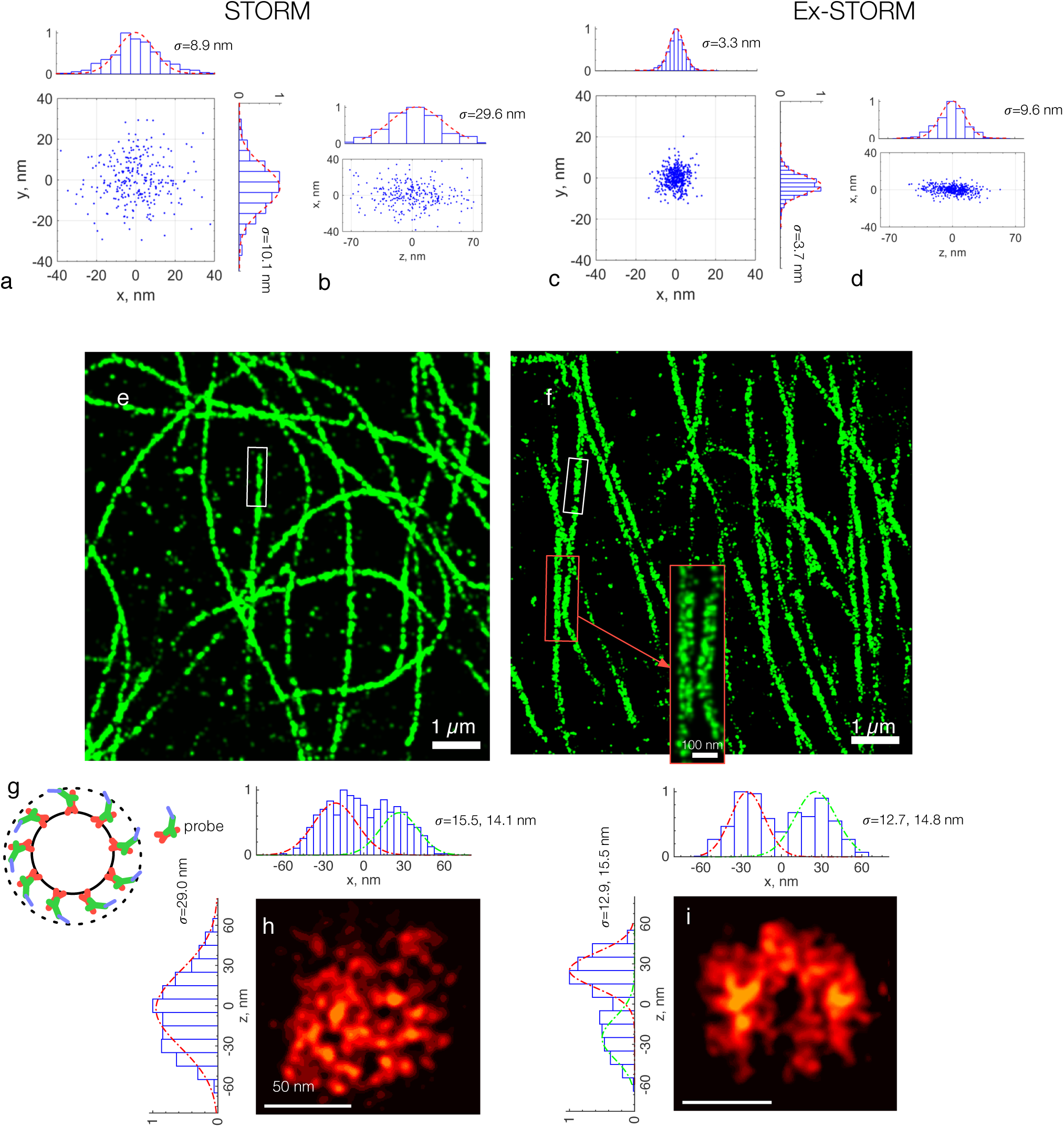
Ex-STORM. (a). Conventional STORM localization precision of Alexa Fluor 647 molecules scattered in fixed U2OS cells are plotted in 2D. Each blinking events are normalized to center at the origin, following ref. 10. The standard deviation *σ* of the distribution in *x* and *y* is 8.9 and 10.1 nm respectively. (b). The distribution of the same localization events in *z-x* plane. The *σ* of the distribution in *z* is 29.6 nm. (c). Localization precision of Alexa Fluor 647 molecules scattered in an expanded gel. The *σ* of the localization events is 3.3, 3.7 and 9.6 nm in *x*, *y*, and *z* (d), respectively. A STORM (e) and an Ex-STORM (f) image of microtubules in U2OS cells. The inset of (f) is a zoom in view of the region selected by the red box in (f). (g) A cartoon illustrates the hollow cross section of microtubule. (h) and (i) are the cross sections of the microtubule selected by the white boxes in (e) and (f), respectively. The histograms in (h) and (i) show profiles of the center sections of the cross sections (−15nm <*x*<15nm for *z*, and −15nm<*z*<15nm for *x*, respectively). The hollow cross section is visible by Ex-STORM (i) but not STORM (h).

The cross-sectional profile of microtubule is commonly used as a standard for evaluating the effective resolution of STORM^1, 10^ (Fig. 2e-f). Microtubule has a hollow cylinder structure (Fig. 2g). When immunofluorescence is used to stain the microtubules, the antibodies bind to the outer side of the cylinder; therefore the cross section of the image of a microtubule should be a hollow circle^10^.

However, such hollow cross section has never been observed before by conventional STORM. While the cross section of a conventional STORM image of a microtubule exhibits a filled circle (Fig. 2h), due to low precision of localization in axial direction, (~69 nm, Fig. 2b), which is larger than the diameter of the microtubule, Fig. 2i shows that hollow cross section can be resolved by Ex-STORM for the first time.

The lateral and axial profiles of an image of microtubule’s cross sections are usually fitted to a double Gaussian^10^; the width of the Gaussian peaks are commonly used as a measure of effective resolution^10, 11^. The improvement of the lateral resolution of Ex-STORM (Fig. 2i, 12.7 and 14.8 nm in *σ*, 29.9 and 34.9 nm in FWHM) is modest from conventional STORM (Fig. 2h, 15.5 and 14.1 in *σ*, 36.5 and 33.2 nm in FWHM); while the improvement of axial resolution is significant, by 2 folds, from 68.3 nm in FWHM to ~33 nm (29.0 nm to 14.2 nm in σ, Fig 2h-i).

The fluorescence probes that we used here are the limiting factor of effective resolution in lateral direction. The point-spread function (PSF) of STORM is convolution of the single molecule localization PSF and the distribution of the probe: *PSF*_STORM_ = *PSF*_Localization_⊗*Probe*. If we assume the localization precision and distribution of the probe are independent from each other, the resolution of STORM is estimated to be: *FWHM*^2^_STORM_ = *FWHM*^2^_Localization_+ *FWHM*^2^_Probe_. Therefore, Ex-STORM achieves the maximum resolution enhancement, when the size of the probe is comparable to or smaller than the localization precision. In this study, we use indirect immunofluorescence method to stain microtubules. The probe consists of a full size primary antibody and full size secondary antibodies; the secondary antibody is conjugated with 12-bp oligonucleotides. The size of the entire probe^10^ is larger than the localization precision in lateral direction (~8 nm), but comparable to the one in axial direction (~23 nm). Therefore, we observes 2 folds enhancement of the effective resolution in axial direction, bringing the *z* resolution on par with *x* and *y*. We expect that the 2.5 folds enhancement of the localization precision can be fully taken advantage of by using sub 10-nm size probes, such as nanobodies or fragment antigen-binding (Fab) fragments.

In summary, we present critical modification to ExM that circumvent the massive bleaching of fluorophores during the polymerization of the polyacrylamide hydrogel. While 50-100% of fluorophores are bleached in the original ExM, bleaching is completely prevented with our methods, which allows imaging of single molecules in expanded samples. We describe efficient sample immobilization method that is compatible to enzymatic oxygen scavenging systems. These improvements clear the barrier between ExM and STORM.

## Materials And Methods

The standard ExM protocol mainly could be found from the ExM website (expansion microscopy.org). We highlight here our improvements to make ExM compatible with STORM.

### 1. DNA Oligonucleotides and Secondary Antibody Conjugation

Oligonucleotide sequence:

5’/5AmMC12/CCGAATACAAAGCATCAACGAAcatctCCGAATACAAAGCATCAACGAA/3′

DNA sequences with 5′ acrydite modification and 3′ amine modifications (Integrated DNA Technologies) were conjugated to the antibodies (Rockland, Affinity Purified goat A-mouse IgG, 6101122) using the Solulink, Antibody Oligonucleotide All-in-One Conjugation Kit (Solulink, A9202001).

### 2. Cultured Cell Preparation and Staining

U2OS cells were cultured on 18x18-1.5 microscope cover glass (FisherScientific, Fisherbrand microscope cover glass, 12541A) in 6-well plates (corning, 3516), in RPMI supplemented with 10%FBS and 1% Penecillin Streptomycin, and grown overnight to 60% confluence in a 37C tissue culture incubator. The following solutions were made in 1x phosphate buffered saline (PBS) and the incubations carried out at room temperature. To preserve microtubule structure, cells were fixed in 3% formaldehyde and 0.1% glutaraldehyde for ten minutes, reduced with 0.1% NaBH_4_ for 7 minutes, and quenched with 100mM glycine for 10 minutes. Cells were permeabilized with 0.2% Triton X-100 for 15 minutes at room temperature, and blocked with 5% Bovine serum albumin (BSA), 10% normal goat serum, and 0.1% Triton X100 for one hour. Samples were incubated with primary antibody (Invitrogen, mouse anti-β-tubulin, 322600) at a concentration of 25μg/ml in the blocking buffer, either 4 hours at room temperature or overnight at 4C. Samples were then incubated with DNA-conjugated secondary antibodies at 25μg/ml in hybridization buffer (2x saline-sodium citrate buffer, 10% dextran sulfate, 1mg/ml yeast tRNA, and 5% goat serum), either 4 hours at room temperature or overnight at 4C.

### 3. Post-Staining With Biotin/Streptavidin

#### 3.1 Biotin conjugated tri-functional label preparation

Oligonucleotide sequence:

5′/Acryd/TTCGTTGATGCTTTGTATTCGGA/3AmMC6T

Complementary tri-functional oligonucleotide were synthesized with a 5′ acrydite and a 3′ amine modifications (Integrated DNA Technologies) and conjugated with NHS-ester biotin (ThermoFisher, EZ-Link NSH-Biotin, 20217) per the manufacturer’s instructions. Streptavidin (New England BioLabs, Streptavidin, N7021S) was conjugated to NHS-ester Alexa647 and NHS-ester Cy3B (GE Life Sciences, Cy3B NHS Ester, PA63100) per the manufacturer’s instructions.

#### 3.2 Biotin oligonucleotide labeling

Samples stained with both primary antibody and secondary antibody conjugated with oligonucleotide (step 2) were incubated with biotin conjugated tri-functional oligonucleotide at a concentration of 10ng/μl in the hybridization buffer overnight at room temperature.

#### 3.3 Polymerization

A monomer solution (1x PBS, 2 M NaCl, 8.63 % sodium acrylate(w/w), 2.5% acrylamide (w/w), 0.15% (w/w) N,N′-methylenebisacrylamide) was mixed with 0.2% (w/w) ammonium persulfate (APS) and 0.2% (w/w) tetramethylethylenediamine (TEMED) to promote polymerization. The monomer solution mixed with the APS and TEMED was then added to samples to a depth of 1mm and incubated for 2 hours at room temperature to allow for the completion of the polymerization process. The size of the gel is measured, which will be used to calculate the expansion ratio later.

#### 3.4 Digestion and dialysis of proteinase K

1mg/ml Proteinase K (Roche, Proteinase K recombinant PCR grade, 03115879001) was added to polymerized samples in ten times volume digestion buffer (50mM Tris pH8, 1mM EDTA, 0.5% Triton X-100, 0.8M guanidine HCl) and incubated at 55C for 20 hours. Digested gels were then placed in 200 excess volumes of streptavidin binding buffer (10mM Tris, 1mM EDTA, 100mM NaCl) to dialyze proteinase K from the gels, and partially expand the gel. Dialysis was repeated with 4 washes of 45 minutes each to ensure complete removal of proteinase K.

#### 3.5 Streptavidin-Alexa647/Cy3B labeling

2x gel volumes of the Alexa647/Cy3B conjugated streptavidin at a concentration of 50μg/ml in streptavidin binding buffer (10mM Tris, 1mM EDTA, 100mM NaCl) was added directly to the gel. Samples were incubated in labeling solution for 24-48 hours at room temperature.

### 4. Post-staining With PNA

An alternative approach to the biotin/streptavidin is by using PNA probes. Multi-color imaging can be achieved by using different PNA sequences.

#### 4.0 List of PNA/ssDNA

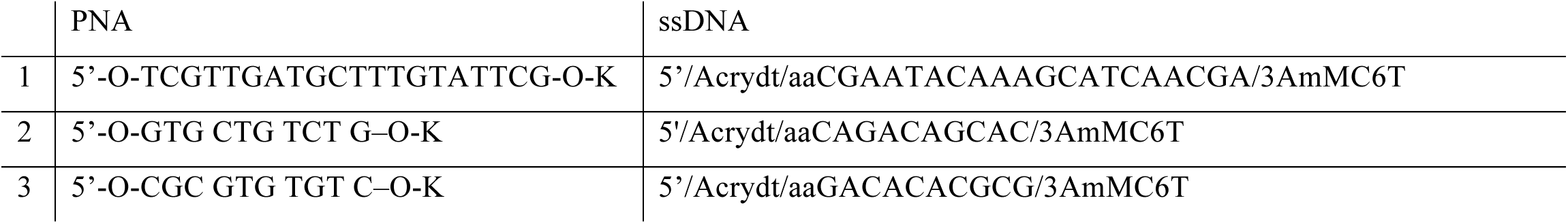

#### 4.1 PNA probe preparation

PNA sequence:

a. 5’-O-TCGTTGATGCTTTGTATTCG-O-K
b. 5’-O-GTG CTG TCT G-O-K
c. 5’-O-CGC GTG TGT C-O-K

Complementary PNA oligonucleotides were synthesized with a 5′ amino and 3′ amine modifications (PNA Bio) and conjugated with NHS-ester Alexa647 (ThermoFisher, AlexaFluor 647 NHS Ester, A37573) and Cy3B (GE Healthcare) per the manufacturer’s instructions.

##### 4.2 DNA oligonucleotides and secondary antibody conjugation

Oligonucleotide sequence:

a. 5′/Acrydt/aaCAGACAGCAC/3AmMC6T
b. 5’/Acrydt/aaCGAATACAAAGCATCAACGA/3AmMC6T
c. 5’/Acrydt/aaGACACACGCG/3AmMC6T

The single-strand DNA oligonucleotides are conjugated to secondary antibodies (Rockland, Affinity Purified goat A-mouse IgG, 6101122) using the Solulink, Antibody Oligonucleotide All-in-One Conjugation Kit (Solulink, A9202001).

##### 4.3 Polymerization

A monomer solution (1x PBS, 2 M NaCl, 8.63 % sodium acrylate(w/w), 2.5% acrylamide (w/w), 0.15% (w/w) N,N′-methylenebisacrylamide) was mixed with 0.2% (w/w) ammonium persulfate (APS) and 0.2% (w/w) tetramethylethylenediamine (TEMED) to promote polymerization. The monomer solution mixed with the APS and TEMED was then added to samples to a depth of 1mm and incubated for 2 hours at room temperature to allow for the completion of the polymerization process.

##### 4.4 Proteinase K digestions

1mg/ml Proteinase K (Roche, Proteinase K recombinant PCR grade, 03115879001) was added to polymerized samples in ten times volume digestion buffer (50mM Tris pH8, 1mM EDTA, 0.5% Triton X-100, 0.8M guanidine HCl, 2.9M NaCl) and incubated at 55C for 20 hours. Digested gels were then placed in 200 excess volumes of 2.9M NaCl solutions to dialyze Proteinase K from the gels. Dialysis was repeated with 4 washes of 45 minutes each to ensure complete removal of proteinase K.

##### 4.5 PNA labeling

Digested and dialyzed gels were incubated with Alexa647 PNA probes. 2x gel volumes of the PNA label at a concentration of 50μg/ml in post-labeling buffer (2x saline-sodium citrate buffer, 1mg/ml yeast tRNA, and 5% goat serum) was added directly to the gel. Samples were incubated in labeling solution for 24-48 hours at room temperature.

#### 5. Expansions and Electrophoresis

Labeled samples were then placed in 500 excess gel volumes of STORM imaging buffer (10mM Tris) to allow for expansion of the gel. Expansion was conducted in imaging buffer with 4 washes of 1hour each. The expanded gel was then cut thin along the gel horizontal axis, to a thickness of approximately 1-2mm, and placed vertically in a precast 0.8% agarose gel with loading lanes 2.5-3mm in thickness. The unbound dye was removed from the gels through electrophoresis, applying 100V for 45-60 minutes. After electrophoresis, the size of the gel was measured again. By comparing the post expansion gel size with the pre-expansion gel size, we determined the expansion ratio.

#### 6. Attachment of Hydrogel to Poly-L-lysine Coated Cover Glass

18x18-#1 cover glass (Fisher, Premium Cove Glass, 12548A) were coated with 500ul of 0.1mg/ml of Poly-lysine (Sigma Aldrich, Poly-L-Lysine solution, 25988630) for 10 miniatures at room temperature. Coverslips were then washed with deionized water and dried overnight in a vacuum chamber. Once the coverslips were dry, expanded and electrophoresies hydrogel samples were placed onto the poly-L-lysine cover glass attaching the hydrogels for imaging.

#### 7. 3D-STORM

3D-*d*STORM imaging of microtubules was carried out on a Nikon Ti inverted microscope equipped with a 60X TIRF objective lens (NA 1.49). An ASI CRISP autofocus system (ASI Imaging) coupled with a 3D piezo stage (Physik Instruments) was used to lock in the focus during imaging. The maximum 639-nm laser (Coherent Genesis, 1W) or a 561nm laser (MPB Communications, 2W) was used followed by activation with 405-nm laser (Coherent OBIS) for Alexa647 and Cy3B stained samples, respectively. A custom Labview (National Instrument) program was used to control the switching between the lasers. A homemade dual-viewer was placed in front of an EMCCD (Andor iXon3, 1024^2^ pixels) for dual color imaging. A 1.5X tube lens was used to achieve a compound magnification of 90X. A 300mm cylindrical lens (Thorlabs) is placed in front of the camera to induce astigmatism for 3D imaging.

We added to the sample an imaging buffer (10mM Tris pH8.0, 10% glucose) composed of 40mM D-glucose, 0.5 mg/ml glucose oxidase (Sigma G6125), 40 μg/ml catalase (Sigma C1345), and 143 mM β-mercaptoethanol (Sigma M6250). Single-molecule blinking events were collected from a FOV of 256×256 pixels with an integration time of 50ms for a total of about 20,000 frames. STORM images were analyzed by ThunderSTORM (GitHub thunderstorm) and MatlabSTORM (https://github.com/ZhuangLab/matlab-storm). Lateral drift was corrected by cross-correlations. Matlab (Mathworks) was also used for data analysis. The images are normalized by the expansion ratio that calculated from the post-and pre-expansion gel size.

## Contributions and Acknowledgements

ZT prepared the samples, collected and analyzed the data. QY set up the STORM microscope and developed the poly-lysine immobilization procedure. JA conjugated the antibodies with DNA oligonucleotides. ZH characterized the expansion ratio of hydrogel in different buffers. HC, ZT and PB designed the experiment. We thank helpful discussions with Professor Fangliang Zhang.

